# Local Microenvironments of capsomer variants in the PBCV-1

**DOI:** 10.1101/2025.01.23.634589

**Authors:** Wenhan Guo, Esther Alarcon, Jason E Sanchez, Chuan Xiao, Lin Li

## Abstract

PBCV-1, a giant virus classified among the Nucleocytoviricota virus (NCV) whose structure has been determined to near atomic resolution. The majority capsomers forming the capsid of PBCV**-**1 are Type I capsomers while five other type of variants have been found in recent high resolution structure. Interestingly, some variants, such as Type V capsomers, are found at particular capsid locations whose roles are unclear. To reveal the roles of a Type V capsomer, we replaced the Type V capsomer by a Type I capsomer to compare the interaction among the two types of capsomer variant, especially the interactions between each of the Type V/Type I capsomer and its local capsid microenvironment. Our results revealed significant differences between Type V and Type I capsomers. Notably, the Type V capsomer demonstrated a stronger binding force to the surrounding capsomers than the Type I capsomer. Moreover, the identified salt bridges between Type V/I capsomers and their surrounding capsomers corroborate the results of electrostatic calculations, further highlighting the important residues involved in these interactions. Understanding these local capsid microenvironments will be essential to elucidate the mechanisms governing viral capsid assembly.

## Introduction

Giant viruses, previously grouped as nucleocytoplasmic large DNA viruses (NCLDVs), are large eukaryotic DNA viruses capable of replicating in both the host cell’s nucleus and cytoplasm [1–3]. Recently, NCLDVs have been officially reclassified into the newly established phylum of *Nucleocytoviricota*. These Nucleocytoviricota viruses (NCVs) have the largest known virions and genomes [4–8], with many genes involved in DNA repair, replication, transcription, and translation [9–13]. NCVs infect various eukaryotic species, including humans [14–18]. Their virions are uniquely structured, typically featuring large, roughly icosahedral-shaped capsids [19–22]. NCVs encompass diverse groups. The discovery of Mimivrus in 1992 propelled the field forward by bringing it to the attention of scientists and leading to the isolation of many other giant viruses [20, 23–26].

Paramecium bursaria Chlorella virus 1 (PBCV-1), one of the earliest and most extensively studied NCVs, contains numerous genes that encode essential functional proteins for the viral life cycle [27–29]. The capsid structure of PBCV-1 has been extensively studied through cryogenic electron microscopy (cryoEM). Early cryoEM studies revealed a quasi-atomic model of the PBCV-1 capsid at 28 Å resolution [30], which was subsequently refined in 2019 to a near-atomic resolution of 3.5 Å for the icosahedrally averaged capsid [19]. More recent subsequent efforts achieved a near atomic resolution of 3.80 Å (PDB ID 8H2I) for the five-fold averaged PBCV-1 capsid structure[31]. These structural analyses revealed unique features of the PBCV-1 capsid, including a specialized portal and other asymmetric components, providing insights into the assembly mechanism of giant virus capsids [32].

PBCV-1’s capsid is composed of a roughly icosahedral outer shell formed by Major Capsid Proteins (MCPs), an inner bilayer lipid membrane, and a thin protein shell between these layers containing minor capsid proteins (mCPs) [19, 21, 31]. The mCPs forms a protein-protein interaction network on the capsid’s inner surface, facilitating its assembly and structural stabilization. The outer capsid of PBCV-1 consists of two distinct types of capsomers: pentameric and pseudo-hexameric, both of which contain jell-roll-fold (JRF), a wedge-shaped β-sandwich fold commonly found in viral capsid proteins. Pentameric capsomers are found exclusively at the vertex of PBCV-1 (as shown in Figure 1A) formed from five single JRF protein. Pseudo-hexameric capsomers, by contrast, are trimers of double JRF MCPs, covering most of the surface of the virus. The capsomers pack closely to cover the icosahedral surface of the virus. Due to differences in height and surface insertions between the two JRFs within the MCPs, the pseudo-hexameric capsomers exhibit a trimeric appearance [20, 33]. These capsomers adopt two distinct orientations, rotated 60° relative to each other, within the capsid’s packed array (as shown in Figure 1B). This orientation variation creates visible grooves on the virus surface, dividing it into distinct patches called symmetrons. The largest patches, known as trisymmetrons, are located on the relatively flat triangular surfaces of the virus.

**Figure 1.**
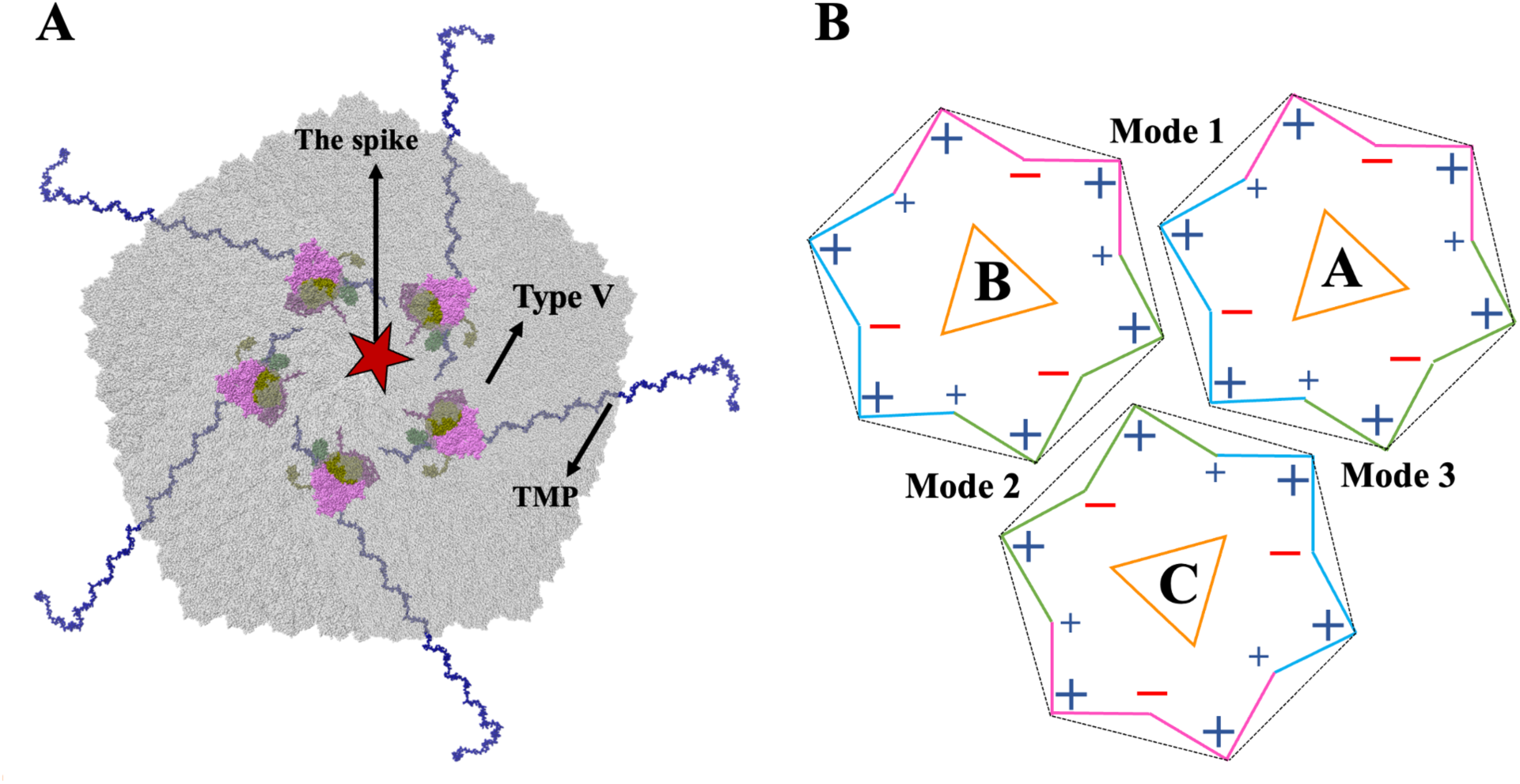
The relative position of the Type V capsomer and capsomer’s three binding modes. (A) The relative position of the Type V capsomer within the outer capsid layer of PBCV-1. (B) Illustration of the three binding modes observed in the outer capsid layer of PBCV-1. Orange triangles indicate capsomers’ orientations.

Our previous study identified three binding modes between capsomers [33]. Mode I, the most prevalent, exists within the trisymmetron and involves capsomers with the same orientation. Modes II and III occur at the boundaries between trisymmetrons, where the capsomers adopt differing orientations. At the pentameric vertices, the orientation differences produce a spiral pattern consisting of three layers of capsomers, collectively termed the pentasymmetron. All NCVs share a conserved pentasymmetron structure, comprising three layers of capsomers (30 MCPs in total) and a penton protein positioned at the five-fold axis. The overall size of a NCV is determined by its trisymmetron size. Interestingly, in both PBCV-1 and Mimivirus, a specialized portal has been identified at one of the 12 five-fold vertices. The finding of unique patterns at the vertices [34] has led to the proposal of a spiral assembly hypothesis, suggesting that capsid assembly initiates at a single vertex (likely the one containing the unique portal) and expands continuously to neighboring vertices [20, 34]. This model aligns with cellular observations [35, 36], which indicate that assembly occurs as a continuous process rather than in a stepwise manner, where patches of symmetrons form first. Notably, large patches of symmetrons have never been observed inside host cells. Additionally, an elongated protein, termed Tape Measure Protein (TMP), was identified running anti-parallelly along the boundaries of trisymmetrons. TMP is hypothesized to play a crucial role in connecting neighboring pentasymmetrons and determining the size of the trisymmetrons [32]. As such, the size of the trisymmetrons directly determines the overall size of the NCVs, highlighting how variations in capsid architecture drive differences in virus size across the group.

The majority of capsomers in NCVs are homotrimers of MCPs. However, heterotrimeric (2:1) capsomers has been proposed first in 2017, based on structural and proteomics data [34]. More recently, the near-atomic, non-icosahedral averaged structure of NCVs convinced the existence of heterotrimeric capsomers by revealing six distinct capsomer variants, including the originally identified, most abundant homotrimeric capsomers. Some of them, Type II, Type III and Type IV are homotrimer like the most abundant Type I MCP. Type I capsomers are the dominant capsomer that cover most surface of the pentasymmetron and the trisymmetron, while Type II and V capsomers are exclusively found in the pentasymmetron, and Type III, IV, and VI capsomers are exclusively found in the trisymmetron [31]. These MCP variants create diverse capsid microenvironments, which likely play critical roles in facilitating the association of fibers, vesicles, and minor capsid proteins. This structural diversity highlights the functional adaptability of NCV capsids, enabling interactions with various components essential for viral assembly and infectivity.

Our previous research utilized computational methods to identify three dominant binding modes (as shown in Figure 1B) within trisymmetrons and at their boundaries, which play a key role in stabilizing the capsid [33, 37]. In this study, we specifically focus on Type V capsomers, composed of three monomers (two MCPv1 and one MCPv2 subunit) and uniquely located near the PBCV-1 capsid’s special vertex, where they associate with a spike portal structure. Using computational approaches, we investigated the local capsid microenvironments influenced by these Type V capsomers. It has been showed that the assembly of PBCV-1 and Mimivirus started from the vertex where the unique portal located. Studying Type V capsomers is therefore particularly significant, as their distinct positioning near the five-fold vertex may offer crucial insights into this continuous assembly mechanism and the initial stages of giant virus capsid formation. Additionally, the Type V capsomer is adjacent to the TMP, suggesting a possible functional interaction between them to extend the assembly from the unique portal vertex to neighboring ones. The elongated TMP, among the intricate network of mCPs beneath the capsomers, extends from one icosahedral fivefold vertex to another. Our previous study reveals the critical role of TMP in the assembly of icosahedral NCV capsids [32]. The close locations of the Type V capsomer and TMP indicate that the Type V capsomer may also contribute to TMP’s function in the structural organization and stability of the viral capsid.

In this work, we employed computational methods to investigate the local capsid microenvironments induced by Type V capsomers in the giant virus PBCV-1. Our multi-scale computational approach combines DelPhi [38, 39], DelPhiForce [40, 41], and NAMD [42] to explore the electrostatic features of Type V pseudo-hexameric capsomers and their surrounding environment. We identified and analyzed the salt bridges that illustrate the interactions among PBCV-1 capsomers to understand their contribution to the overall electrostatic balance of the protein. To assess the significance of Type V pseudo-hexameric capsomers, we also performed calculations using Type I pseudo-hexameric capsomers to compare their effects on the local capsid microenvironments. Our findings reveal significant differences in the electrostatic properties between Type V and Type I pseudo-hexameric capsomers. Specifically, Type V capsomers exhibit stronger attractive forces towards surrounding capsomers compared to Type I capsomers. The non-uniform distribution of Type V capsomers near this unique vertex may play a pivotal role in the assembly of PBCV-1.

## Methods

### Complex Structure Preparation and Molecular Dynamics simulations

The Type V pseudo-hexameric capsomer and its surrounding capsomers were extracted from PBCV-1 (PDB ID 8H2I [31]), a near-atomic resolution, non-icosahedrally averaged model of PBCV-1 obtained via cryo-EM at a resolution of 3.80 Å. To evaluate the impact of local capsid microenvironments created by Type V pseudo-hexameric capsomers, we also constructed a model replacing the Type V capsomer with a Type I pseudo-hexameric capsomer for comparative analysis. After obtaining the complex structures of both Type V and Type I capsomers with surrounding capsomers, structural clashes were resolved using CHARMM-GUI [43], a tool specifically designed for preparing molecular structures. Once the structures were free of clashes, a 20 ns Molecular Dynamics (MD) simulation was performed for each complex using NAMD 2.12 [42] to optimize their conformations. The MD simulations were conducted at a constant temperature of 300 K, maintained by a Langevin thermostat with a damping coefficient of 1/ps. The pressure was kept at 1 atm using a Nosé–Hoover Langevin piston barostat, which operated with a decay period of 50 fs. Periodic boundary conditions were employed throughout the simulations to simulate an infinite system. The CHARMM36m force field [44] was applied to model the protein dynamics, and a 150 mM KCl concentration was used to ionize the system, reflecting physiological conditions. Prior to running the simulations, the system underwent a 20,000-step energy minimization to eliminate any unfavorable interactions and bring the system closer to equilibrium. During the MD simulations, residues with any atom within 10 Å of the Type V/Type I capsomer and surrounding capsomers at the binding interfaces were designated as interfacial residues. These residues were left unconstrained, allowing them to move freely during the simulation, while non-interfacial residues were constrained to focus the dynamics on the key binding interactions. The simulations were visualized using VMD [45], and the Root Mean Square Deviation (RMSD) was calculated for each frame to monitor the structural stability of the complexes over time. Based on the RMSD plot (Figure S1), which showed stable structures after initial fluctuations, the last 10 ns of the simulations were selected for further analysis, as this period demonstrated structural stability. Additionally, the last frame (4000th frame) from each MD simulation was extracted to use in subsequent calculations, since the protein complex structures were well-equilibrated by the end of the simulation. The complex structures of both Type V and Type I capsomers with their surrounding capsomers, as used for further electrostatic and force calculations, are depicted in Figure 2. In the visualizations, the capsomer monomer, MCPv1 monomers, and MCPv2 monomers are shown in green, purple, and grey, respectively. The color scheme was used consistently throughout the work. To improve the clarity of these visualizations, the long tail domain (residues 2-67) and the upper region of the Type V capsomer were removed for electrostatic potential and electric field line calculations. This selective removal of structural components was essential for focusing the analyses on the key interaction sites between the capsomers (MCPs). This detailed preparation and simulation process allowed us to accurately model the local capsid microenvironments around the Type V/Type I capsomers, providing insights into how structural differences between capsomers influence the stability and interactions within the viral capsid.

**Figure 2.**
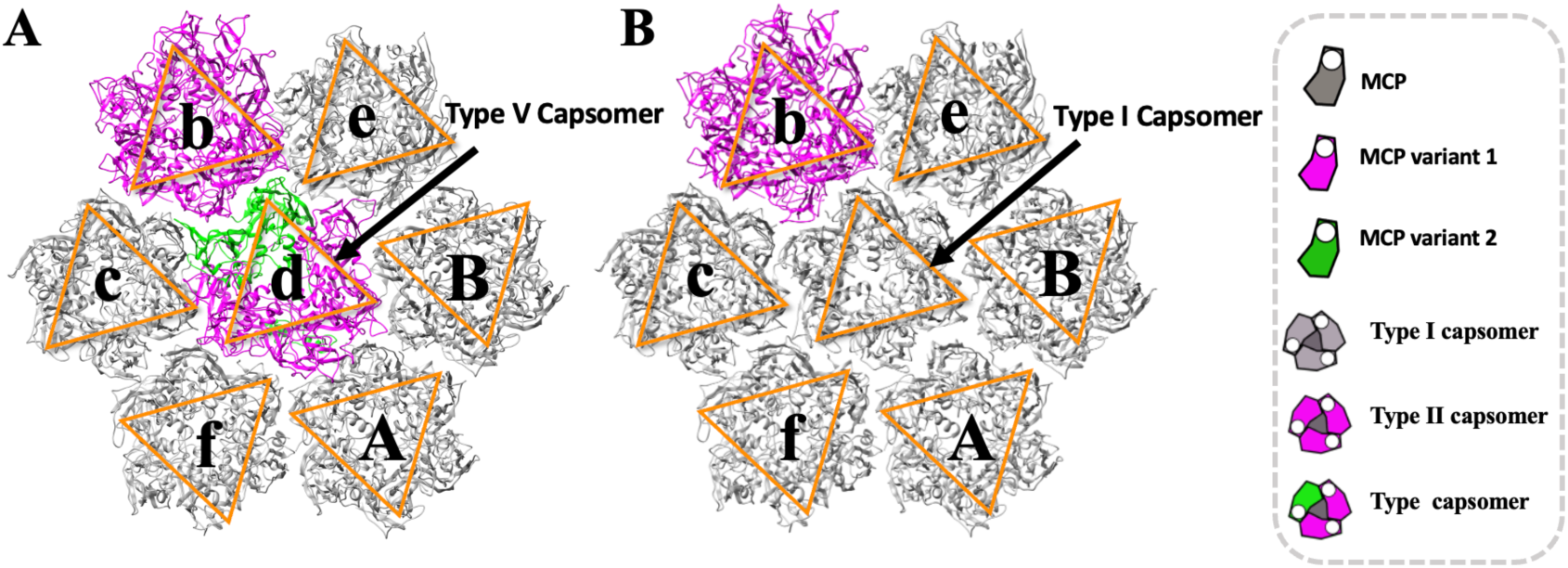
Complex structures of Type V/Type I capsomer with surrounding capsomers. (A) Complex structure of the Type V capsomer with surrounding capsomers. (B) Complex structure of the Type I capsomer with surrounding capsomers. Each capsomer is labeled with letters according to Shao’s work [31]. Capsomers are labeled by orange triangles to show their orientations.

### Electrostatic potential

To investigate the local capsid microenvironments created by interactions between the Type V/Type I capsomer and surrounding capsomers, DelPhi [38, 39] was used to calculate the electrostatic potential. DelPhi is a widely used computational tool for computing the electrostatic potential of biomolecules [46–51]. It solves the Poisson-Boltzmann equation (PBE) using the finite difference method, as shown below:

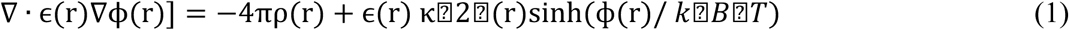

In this equation, ɛ r represents the dielectric permittivity, ф r displays the electrostatic potential, ρ r is the permanent charge density, κ is the Debye-Hckel parameter, *k B* is the Boltzmann constant, and *T* is temperature. The PBE allows for the calculation of the electrostatic potential in and around biomolecules, considering both solvent effects and ionic strength, which are crucial for understanding interactions within a complex biological system. When applying DelPhi to calculate the electrostatic potential, the protein filling percentage within the calculation box was set to 70%, which balances computational efficiency with accurate representation of the molecular environment. The probe radius for the molecular surface was set to 1.4 Å, simulating the size of a water molecule. A salt concentration of 150 mM was applied to mimic physiological conditions, which is critical in determining the ionic strength of the system. The dipolar boundary condition was used to ensure an accurate representation of the electrostatic potential far from the molecular surface, thereby preventing artificial boundary effects. DelPhi generated three-dimensional electrostatic potential maps of the interactions between the Type V/Type I capsomer and the surrounding capsomers. These maps provided a visual representation of the electrostatic environments surrounding the capsomers, highlighting regions of positive and negative charge distribution. The electrostatic potential maps were visualized using UCSF Chimera [52], a molecular visualization tool that allowed for detailed analysis of the potential surfaces. The color scale for the maps was set between -1.0 and 1.0 kT/e, where negative values (in red) indicate areas of negative electrostatic potential and positive values (in blue) represent areas of positive potential. This visualization method provided an intuitive understanding of how charge distributions vary across different regions of the capsid. The calculated electrostatic potentials revealed key differences between the Type V and Type I capsomer interactions with surrounding capsomers. Specifically, the non-uniform charge distribution of Type V capsomer created distinct electrostatic microenvironments, which likely play a crucial role in viral capsid assembly. Areas of high positive or negative potential may serve as hotspots for protein-protein interactions, guiding the alignment and binding of capsomers during assembly. Comparing the electrostatic potentials of Type V and Type I capsomer interactions further illustrated how variations in charge distribution could influence the stability and strength of these interactions. By providing a detailed, visual representation of the electrostatic landscapes within the capsid, this analysis offers valuable insights into the molecular forces driving viral capsid assembly. The observed differences in electrostatic potential between the two types of capsomers may help explain the functional roles of these capsomers in maintaining structural integrity and promoting the assembly of the PBCV-1 capsid.

### Electric Field Lines

To gain further insight into the detailed interactions between the Type V/Type I capsomer and surrounding capsomers, electric field lines were calculated using DelPhi [38, 39] and visualized with VMD [45]. The electric field lines reveal the directional forces between the Type V/Type I capsomer and its surrounding capsomers in the complex structures. The color scale was set between -1.0 and 1.0 kT/e, corresponding to negative and positive electrostatic potentials, respectively. To enhance visualization clarity, the surrounding capsomers were positioned 20 Å away from the Type V/Type I capsomer. This separation allowed a clearer observation of the electrostatic field lines, highlighting regions with stronger electrostatic interactions. The density of these field lines reflects the magnitude of the electrostatic forces, with denser lines indicating stronger interactions between the Type V/Type I capsomer and the surrounding capsomers. The visualization of the electric field lines offers key insights into the electrostatic microenvironment of the capsid. In particular, it helps to understand how the non-uniform charge distribution of Type V capsomer influences the interaction patterns with its surrounding capsomers, potentially playing a critical role in the viral capsid assembly process. By comparing Type V and Type I capsomer interactions, these visualizations allow us to examine how differences in charge distribution contribute to distinct electrostatic environments within the viral capsid.

### Electrostatic Force

To quantify the electrostatic interactions between the Type V/Type I capsomer and surrounding capsomers, we used DelPhiForce [40, 41], a computational tool designed to calculate electrostatic forces by solving the Poisson-Boltzmann equation (PBE), as described in Equation (1). The surrounding capsomers were systematically separated from the Type V/Type I capsomer by distances ranging from 6 Å to 40 Å, with 4 Å increments, using StructureMan [53]. This incremental separation allowed for a detailed analysis of how electrostatic forces change with distance between the capsomers. The directions of the electrostatic forces between the Type V/Type I capsomer and surrounding capsomers were visualized as blue arrows using Visual Molecular Dynamics (VMD) [45]. These arrows depict not only the direction of the forces but also provide an intuitive understanding of the attractive or repulsive nature of the interactions at different distances. The magnitude of these electrostatic binding forces was also calculated at each distance step, offering quantitative insights into the strength of the interactions. To further illustrate the relationship between the Type V/Type I capsomer and its surrounding capsomers, we plotted the electrostatic binding forces as a function of distance using histograms. These histograms allowed us to observe how the strength of the interaction changes as the capsomers move farther apart, highlighting the differences in electrostatic behavior between Type V and Type I capsomers. Notably, the analysis revealed that Type V capsomer exhibited stronger attractive forces over a wider range of distances compared to Type I capsomer. This suggests that the unique charge distribution of Type V capsomer may contribute to a more robust electrostatic network within the viral capsid, which could be a key factor in the assembly process of PBCV-1. By combining electric field line visualizations and quantitative electrostatic force measurements, we have provided a comprehensive analysis of the electrostatic microenvironments around the Type V/Type I capsomer and their interactions with surrounding capsomers. These findings offer valuable insights into how the local microenvironment electrostatic properties of specific capsomers influence viral capsid assembly and stability.

### Salt bridges

Salt bridges can provide important insights into the stability and strength of protein-protein interactions within the viral capsid. To investigate the role of interfacial residues that significantly contribute to the binding interactions between the Type V/Type I capsomer and surrounding capsomers, salt bridges were analyzed based on the results of the molecular dynamics (MD) simulations. The last 10 ns (2000 frames) of the MD simulations were selected for this analysis, as the structures had reached equilibrium by this point. Salt bridges were identified using the salt bridge tool in VMD, with a distance threshold set to 4 Å, which is commonly used to define a salt bridge interaction. This threshold ensures that only relevant, strong ionic interactions between charged residues at the binding interface are considered. The percent occupancy of each salt bridge during the MD simulations was calculated, allowing us to assess the stability and prevalence of each interaction over time. Salt bridges with a percent occupancy of less than 10% during the simulations were excluded from the analysis, as these are likely to be transient and have minimal impact on the overall stability of the capsomer-capsomer interaction. Conversely, salt bridges with high occupancy were considered significant contributors to the binding interactions. These stable interactions are expected to play a critical role in maintaining the structural integrity of the capsid, especially in the context of the non-uniform distribution of charges observed in Type V capsomer.

The presence of strong, stable salt bridges may be crucial for facilitating capsid assembly, as they provide both structural stability and specific electrostatic interactions required for the alignment and binding of capsomers. By identifying and quantifying these salt bridges, we could gain valuable insights into the molecular mechanisms underlying the assembly and stability of the PBCV-1 capsid. This analysis highlights the importance of interfacial residues and their contributions to the binding strength between capsomers.

## Results and Discussions

To investigate the roles of Type V capsomers and the local capsid microenvironments induced by them in the giant virus PBCV-1, we compared the electrostatic features of the interactions of the Type V/Type I capsomers and their surrounding capsomers.

### Electrostatic potential comparison

Electrostatic features play a crucial role in protein-protein interactions [46, 54–56], including the assembly of the capsid. Our calculations show that the net charge of Type I capsomer is neutral. On the other hand, the Type V capsomer is significantly negatively charged (-16e). Such a significant charge differences leads to distinct binding mechanisms with their surrounding capsomers.

Furthermore, the electrostatic potentials on the surfaces of the Type V and Type I capsomers were calculated and shown in Figure 3. Structures of Type V and I with their surrounding capsomers are shown in figure 3A and 3B. In the top (looking from outside of capsid) view of the Type V capsomer (Figure 3C), the structure is predominantly negatively charged, indicating a strong overall negative electrostatic environment on capsomere’s outside surface. In contrast, the top view (looking from outside of capsid) of the Type I capsomer (Figure 3D) reveals a more mixed distribution of negative, positive, and neutral charges across the surface, creating a more heterogeneous electrostatic landscape. When examining the bottom (looking from inside of the capsid) view of the Type V capsomer (Figure 3E), the structure shows a predominantly positively charged central region, surrounded by smaller, negatively charged areas. In comparison, the bottom view of the Type I capsomer (Figure 3F) is even more positively charged than the bottom view of the Type V capsomer. This difference can be attributed to the distinct composition of the capsomers: the Type V capsomer is made up of two MCPv1 subunits and one MCPv2 subunit, while the Type I capsomer consists entirely of three MCPv2 subunits. This compositional difference leads to significantly different electrostatic potentials between the two capsomers, resulting in distinct electrostatic interactions with the surrounding capsomers. The marked differences in the electrostatic potentials of the Type V and Type I capsomers are critical for understanding their roles in viral capsid assembly. These differences suggest that the Type V capsomer may have a unique role near the viral capsid’s vertex, where its specific charge distribution could be essential for attracting or repelling surrounding capsomers. The presence of a Type V capsomer near the unique vertex rather than a Type I capsomer, could be integral to initiating the assembly process of the viral capsid.

**Figure 3.**
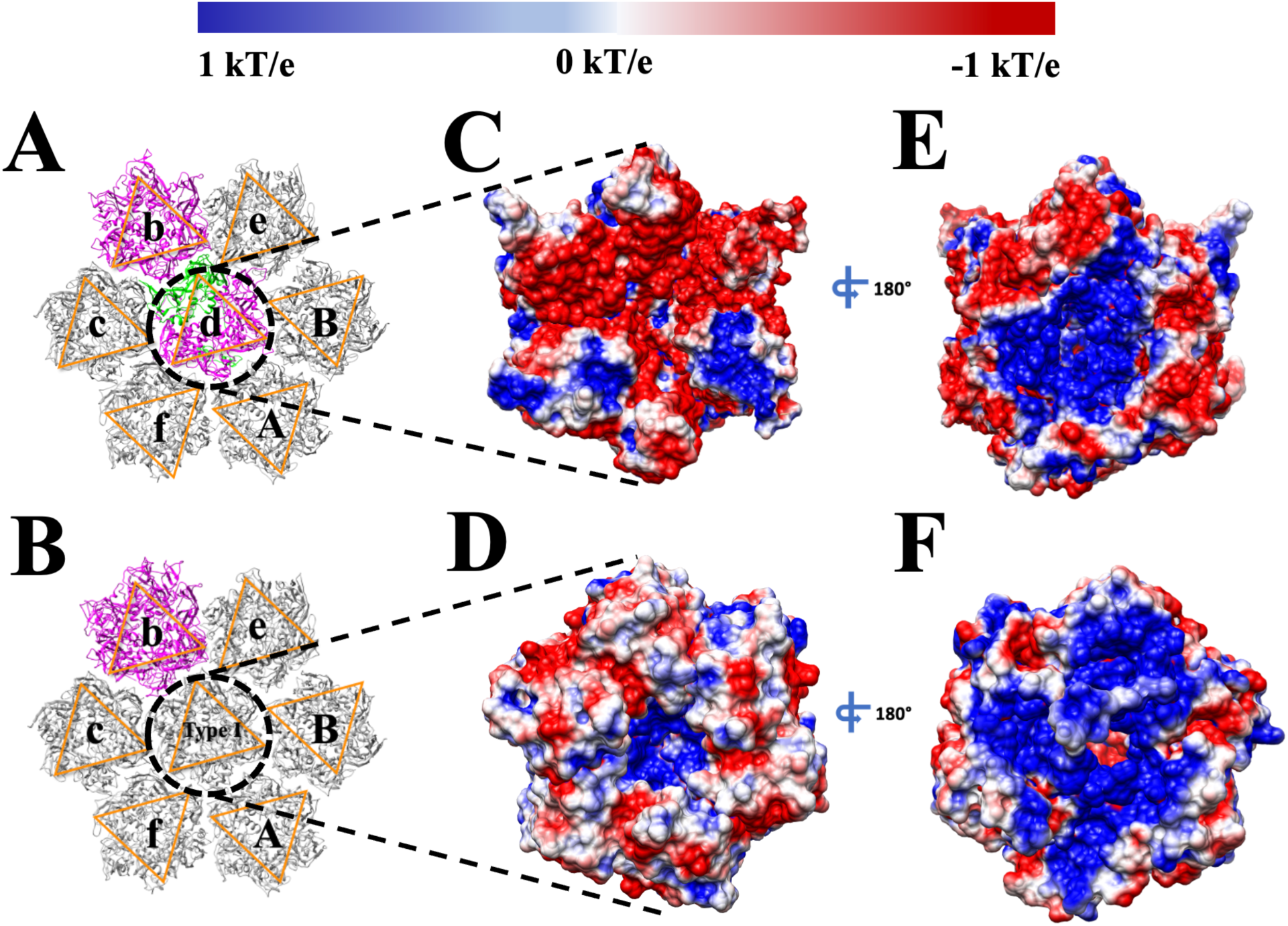
Structures and electrostatic potentials of Type Ⅴ/ Type I capsomer. (A) Structures of Type Ⅴ capsomer with surrounding capsomers. (B) Structures of Type I capsomer with surrounding capsomers. (C) Top (looking from outside of capsid) view of electrostatic potentials of Type Ⅴ capsomer with surrounding capsomers. (D) Top (looking from outside of capsid) view of electrostatic potentials of Type I capsomer with surrounding capsomers. (E) Bottom (looking from inside of the capsid) view of electrostatic potentials of Type Ⅴ capsomer with surrounding capsomers. (F) Bottom (looking from inside of the capsid) view of electrostatic potentials of Type I capsomer with surrounding capsomers. MCP monomer variants are displayed in the same colors as shown in Figure 2. The color scale range for electrostatic potential is from -1.0 to 1.0 kT/e. Capsomers are labeled by orange triangles to show their orientations.

The electrostatic potentials at the binding interfaces of the Type V/Type I with the surrounding capsomers are shown in Figure 4. To better visualize the interaction interfaces, the surrounding capsomers and the Type V/Type I capsomers were each rotated 90 degrees in opposite directions along the center of mass of the Type V/Type I capsomers with surrounding capsomers. This rotation revealed the interfacial electrostatic potentials between the Type V/Type I capsomers and their corresponding surrounding capsomers. The interactions between the Type V/Type I capsomer and their surrounding capsomers are driven by the net charges at their interfaces: opposite charges lead to attractive interactions, while like charges result in repulsion. The difference in charge between the two capsomers directly influences their conformational interactions with the surrounding capsomers. Specifically, the changes in electrostatic potential at the interfaces of the surrounding capsomers are a result of the structural differences between the Type V and Type I capsomers.

**Figure 4.**
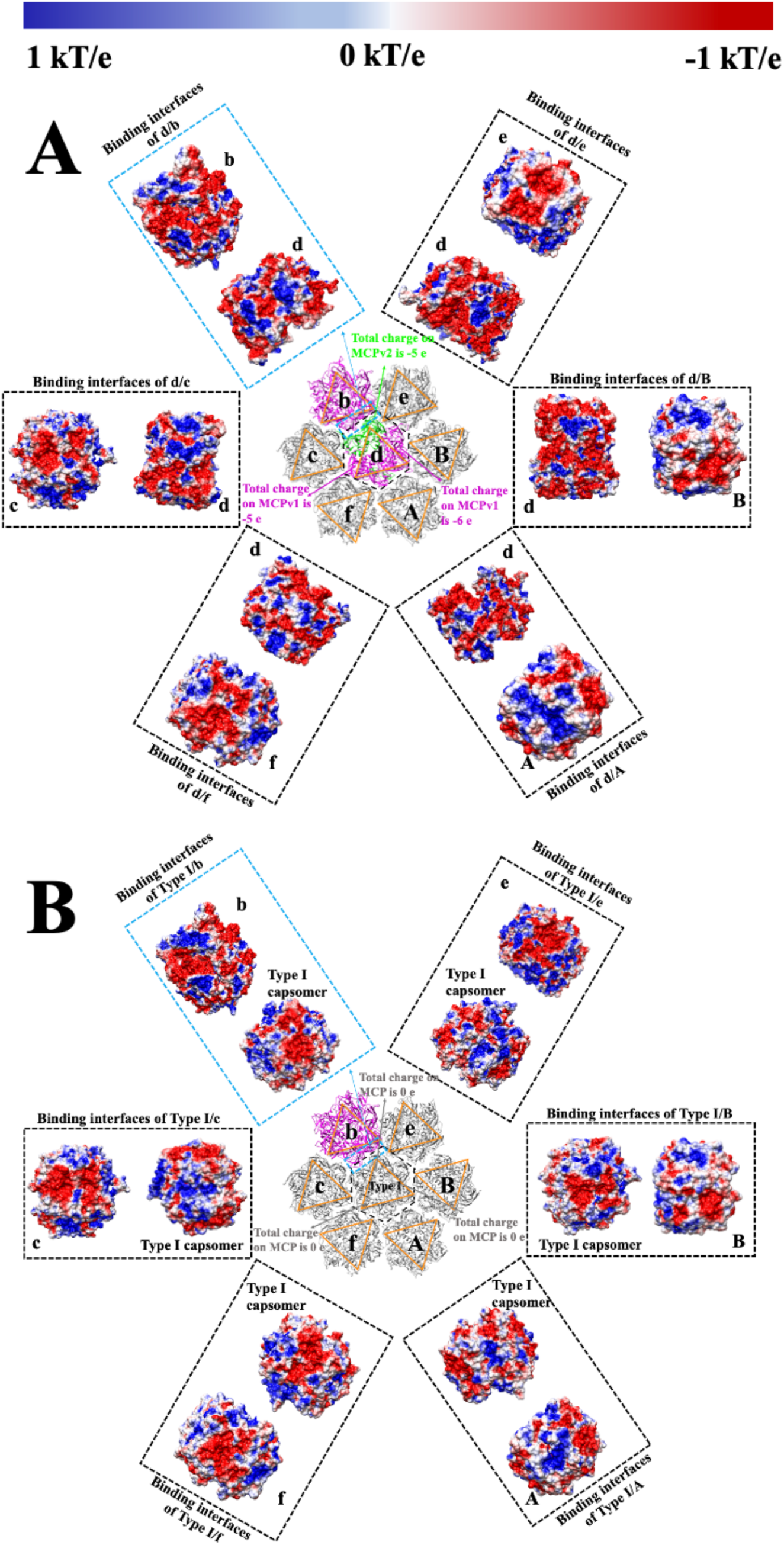
Electrostatic potentials at the binding interfaces of Type Ⅴ/ Type I capsomer with surrounding capsomers. (A) Surrounding capsomers’ interfacial electrostatic potentials with Type Ⅴ capsomer. Surrounding capsomers and the Type V capsomer were each rotated 90 degrees in opposite directions along the center of mass of the Type V capsomer with surrounding capsomers. (B) Surrounding capsomers’ interfacial electrostatic potentials with Type I capsomer. Surrounding capsomers and the Type I capsomer were each rotated 90 degrees in opposite directions along the center of mass of the Type I capsomer with surrounding capsomers. MCP monomer variants are displayed in the same colors as shown in Figure 2. The total charge on these single monomers was marked. The color scale range for electrostatic potential is from -1.0 to 1.0 kT/e. Capsomers are labeled by orange triangles to show their orientations.

Figure 4 shows that the electrostatic potential at the binding interfaces between the surrounding capsomers and the central capsomer differs only slightly. This similarity is expected, as the surrounding capsomers are the same before undergoing MD simulations. However, the electrostatic potential at the binding interface between the central Type V/Type I capsomer and the surrounding capsomers is entirely different. Consequently, the binding regions and sites between the Type V/Type I capsomer and the surrounding capsomers are also totally different. From the electrostatic potential results, the Type V capsomer shows larger opposite-charged interaction regions with surrounding capsomers compared to the Type I capsomer. The difference results in the stability change in the capsid assembly. The detailed electrostatic features, including electric field lines and electrostatic forces, are discussed in the following sections.

### Electric Field Lines Comparison

To explore the electrostatic interactions between the Type V/Type I capsomers and their surrounding capsomers, we calculated the electric field lines for the two complexes, as shown in Figure 5. The electric field lines provide visual insight into the strength and direction of electrostatic forces between the capsomers and their neighboring capsomers. To enhance visualization, the surrounding capsomers were separated from the Type V/Type I capsomers by 20 Å, allowing a clearer view of the interaction regions. The density of the electric field lines directly correlates with the strength of the electrostatic interactions: denser lines indicate stronger forces, while sparser lines reflect weaker interactions. In the complex with the Type V capsomer at the center (Figures 5AB), the electric field lines between the Type V capsomer and the surrounding capsomers are notably denser than the Type I capsomer in the center (Figure 5CD). This suggests strong electrostatic attractions between the Type V capsomer and the surrounding capsomers, reinforcing the idea that the charged Type V capsomer plays a critical role in binding and stabilizing the viral capsid structure. In contrast, when the Type I capsomer is at the center of the complex (Figures 4CD), the electric field lines between the Type I capsomer and surrounding capsomers are much sparser. It indicates significantly weaker electrostatic interactions compared to the Type V capsomer complex. Consequently, the sparser electric field lines indicate weaker electrostatic forces between the Type I capsomer and surrounding capsomers, which may result in a less stable assembly. The stark difference in electric field line density between the two complexes supports the hypothesis that the charged Type V capsomer plays a pivotal role in the assembly process of PBCV-1. Its stronger electrostatic interactions with surrounding capsomers likely enhance the stability and integrity of the capsid, especially near the unique vertex where the Type V capsomer is located.

**Figure 5.**
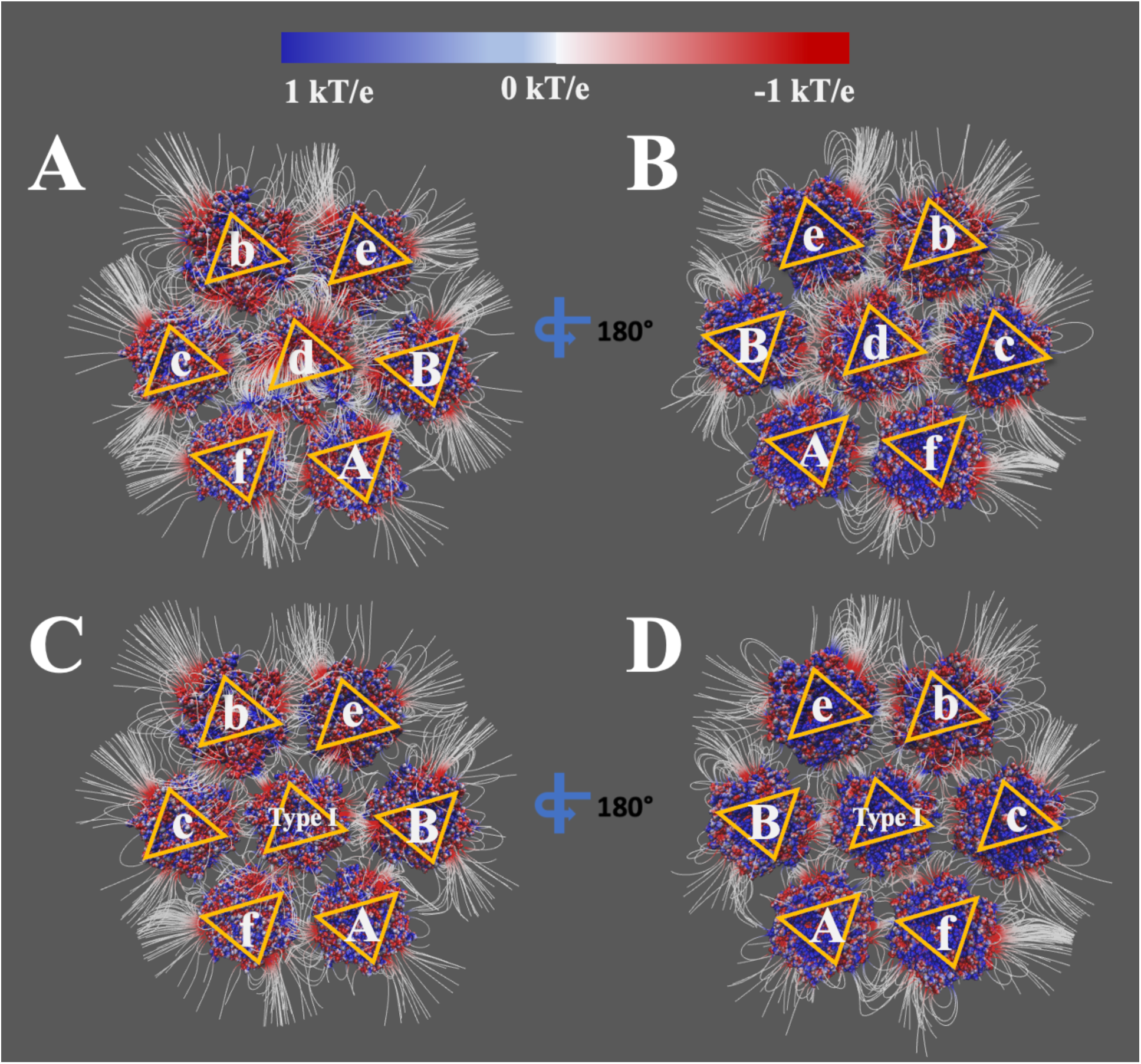
Electric field lines at the binding interfaces of Type Ⅴ/ Type I capsomer with surrounding capsomers. (A) (B) are the top (looking from outside of capsid) view and the bottom (looking from inside of the capsid) view of the electric field lines between the Type Ⅴ capsomer with surrounding capsomers, respectively. (C) (D) are the top (looking from outside of capsid) view and the bottom (looking from inside of the capsid) view of the electric field lines between the Type I capsomer with surrounding capsomers, respectively. Capsomers are labeled by orange triangles to show their orientations.

In summary, the electric field line comparison between the Type V and Type I capsomers highlights the importance of electrostatic interactions in the assembly and stability of the PBCV-1 capsid. The stronger electrostatic forces observed with the Type V capsomer, as indicated by the denser field lines, suggest that this charged capsomer is essential for forming stable interactions with surrounding capsomers, thereby playing a crucial role in the viral assembly process. The following sections will further explore these interactions through quantitative analysis of electrostatic forces and salt-bridges.

### Electrostatic Forces Comparison

DelPhiForce [40, 41] was employed to calculate the electrostatic forces involved in the interactions between the Type V/Type I capsomers and their surrounding capsomers. To compare the differences between the two complexes, we systematically separated the surrounding capsomers from the Type V/Type I capsomers at distances ranging from 6 Å to 40 Å, with increments of 2 Å, using StructureMan [53]. At each separation distance, the electrostatic forces between the capsomers and their surrounding capsomers were calculated using DelPhiForce. These calculations allowed us to analyze the directions and strengths of the electrostatic forces at varying distances (Figure 6).

**Figure 6.**
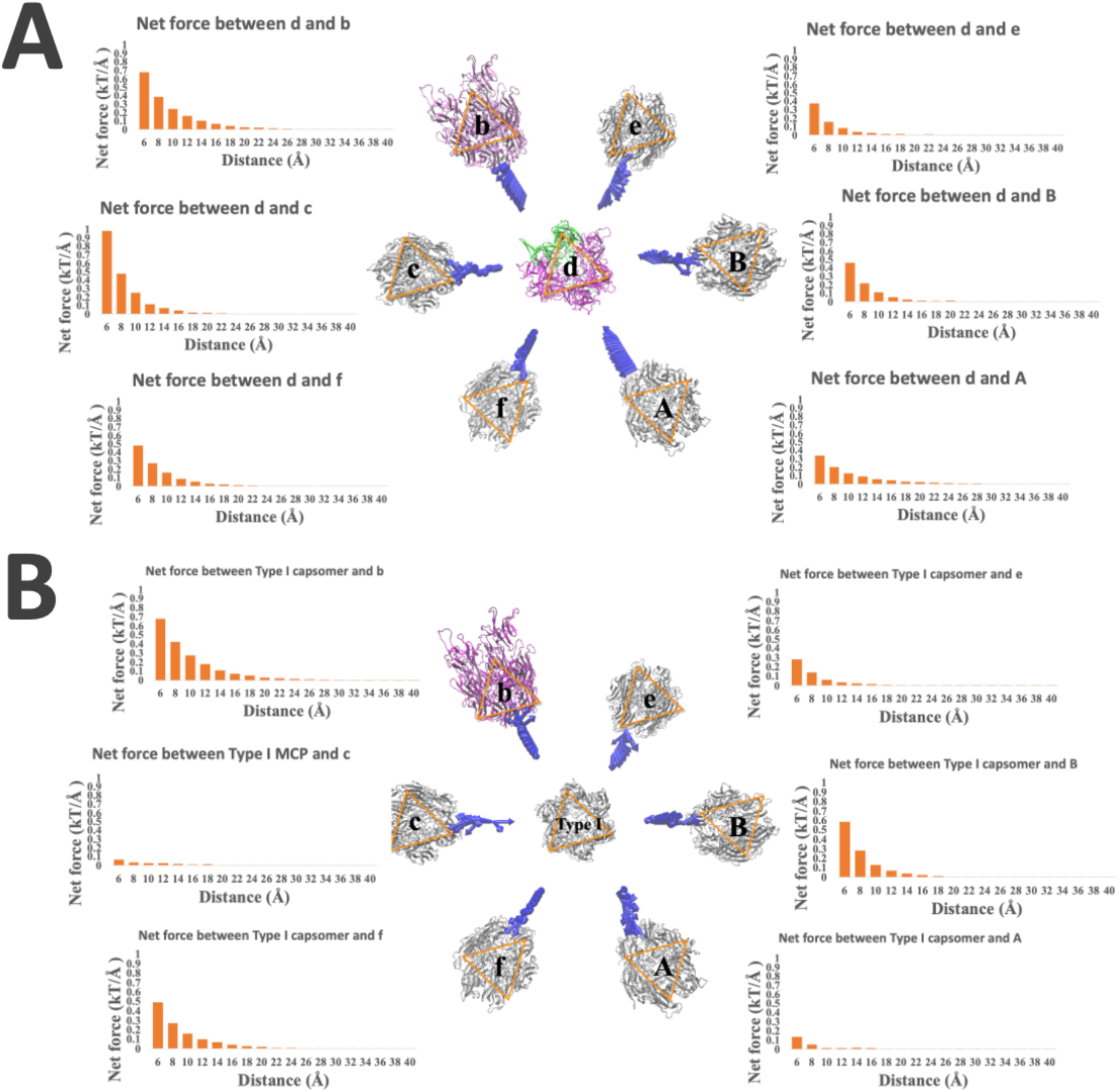
Electrostatic forces between Type Ⅴ/ Type I capsomer with surrounding capsomers. (A) shows the directions and magnitude of the electrostatic forces felt by the Type Ⅴ at distances from 6 Å to 40 Å with a step size of 2 Å from surrounding capsomers. (B) shows the directions and magnitude of the electrostatic forces felt by the Type I at distances from 6 Å to 40 Å with a step size of 2 Å from surrounding capsomers. Capsomers are labeled by orange triangles to show their orientations.

The directions of the net electrostatic forces between the Type V/Type I capsomers and their surrounding capsomers are visualized as blue arrows in Figure 6. To facilitate comparison, the arrows have been normalized to the same size for clarity. In the complex with the Type V capsomer at the center (Figure 6A), the arrows indicate that the electrostatic forces are generally directed toward the Type V capsomer, demonstrating strong attractive forces between the capsomer and all surrounding capsomers. This consistent attraction suggests that the Type V capsomer exerts a strong stabilizing influence on the neighboring capsomers, pulling them toward its center. Among these interactions, d-c has the strongest electrostatic force (0.98 kT/Å at 6 Å), followed by d-b (0.68 kT/Å at 6 Å). In contrast, the complex with the Type I capsomer at the center (Figure 6B) shows more varied electrostatic interactions. The B and e capsomers exhibit attractive forces toward the Type I capsomer, while the b and f capsomers display weaker sliding forces, as indicated by the direction of the arrows. These sliding forces suggest weaker, more lateral interactions rather than direct attraction. Notably, the A capsomer exhibits repulsive forces, as the arrows point away from the Type I capsomer, indicating that this capsomer is pushed away rather than pulled toward the center. The c capsomer initially shows attraction to the Type I capsomer at close range (6-8 Å), but as the distance increases, the direction of the electrostatic force shifts, indicating repulsive forces beyond 8 Å. This diverse array of interactions in the Type I complex contrasts sharply with the uniform attraction observed in the Type V complex. In conclusion, the Type V capsomer demonstrates stronger and more consistent electrostatic forces toward surrounding capsomers compared to the Type I capsomer. The consistent attractive forces observed in the Type V complex suggest a more stable and cohesive interaction network, which may play a critical role in the assembly and stability of the PBCV-1 capsid.

In addition to the directional analysis, we also quantified the magnitudes of the electrostatic forces between the Type V/Type I capsomers and their surrounding capsomers, as shown in Figure 6. As expected, the electrostatic forces decrease with increasing distance between the capsomer and surrounding capsomers, following Coulomb’s law. However, the magnitudes of the forces in the Type V capsomer complex are consistently stronger than those in the Type I capsomer complex. This finding reinforces the idea that the charged Type V capsomer exerts more substantial forces on its surrounding capsomers compared to the neutral Type I capsomer. In the Type V capsomer complex, the attractive forces range from 0.34 to 0.98 kT/Å, indicating a strong electrostatic pull between the capsomer and surrounding capsomers. In contrast, the Type I capsomer complex exhibits a more varied force landscape, with two capsomers experiencing attractive forces, two showing sliding forces, and two displaying weak repulsive forces. This difference in electrostatic force distribution further emphasizes the distinct roles played by the Type V and Type I capsomers in capsid assembly. Overall, the Type V capsomer consistently shows stronger electrostatic forces, both in terms of direction and magnitude, compared to the Type I capsomer. This finding aligns with the earlier analyses of electrostatic potentials and electric field lines, reinforcing the conclusion that the Type V capsomer plays a critical role in stabilizing the local microenvironment and facilitating the assembly of the PBCV-1 capsid.

Moreover, our results on electrostatic forces are consistent with our previous findings on PBCV-1 [33]. The previous study found three binding modes among capsomers based on their orientations inner and inter trisymmetrons. In that study, binding Mode I consists of two capsomers aligned in the same orientation within the trisymmetron structures, showing attractive electrostatic interactions. Each of the binding Mode II and Mode III exists at the boundary between symmetrons with two capsomers oriented at a 60-degree difference. Binding Mode I shows attractive electrostatic interactions, while binding Mode III results in repulsive electrostatic interactions.

However, because that work focus was on binding modes inner and inter the trisymmetron structures, it primarily considered interactions among Type I capsomers. In contrast, this research examines capsomers located near the unique vertex, including three types: Type V (d), Type II (b), and Type I capsomers (c, f, A, B, e). Thus, in this work, capsomers ‘d,’ ‘Type I,’ ‘c,’ ‘b,’ and ‘e’ share the same orientation, while capsomers ‘B,’ ‘A,’ and ‘f’ differ by 60 degrees. Consequently, our electrostatic force analysis shows the interactions of d-c, d-b, d-e, Type I-c (within 8 Å), Type I-b, and Type I-e belong to binding Mode 1 which result in attractive electrostatic interactions. The interaction of d-f, d-B, Type I-f, and Type I-B belong to binding Mode II showing attractive electrostatic interactions. The interaction between Type I and A is in binding Mode III, therefore shows repulsive electrostatic interactions.

### Salt Bridges Comparison

Salt bridges between the Type V/Type I capsomers and their surrounding capsomers were analyzed based on the results of the MD simulations. Salt bridges with occupancies above 10% were considered significant and are shown in Figure 7. The results reveal a remarkable difference in the number and stability of salt bridges formed between the two capsomer types and their respective surrounding capsomers. Overall, the number of salt bridges formed between the Type V capsomer and surrounding capsomers is substantially higher than those formed with the Type I capsomer. It suggests that the Type V capsomer establishes more extensive and stable interactions with its neighboring capsomers, which may be crucial for maintaining the structural integrity of the viral capsid. In contrast, no salt bridges are formed with half of the surrounding capsomers when the Type I capsomer is in the center, indicating a lack of interaction at these interfaces. This absence of salt bridges highlights the limited ability of the Type I capsomer to engage in stabilizing interactions, suggesting it may be less effective in capsid assembly compared to the Type V capsomer.

**Figure 7.**
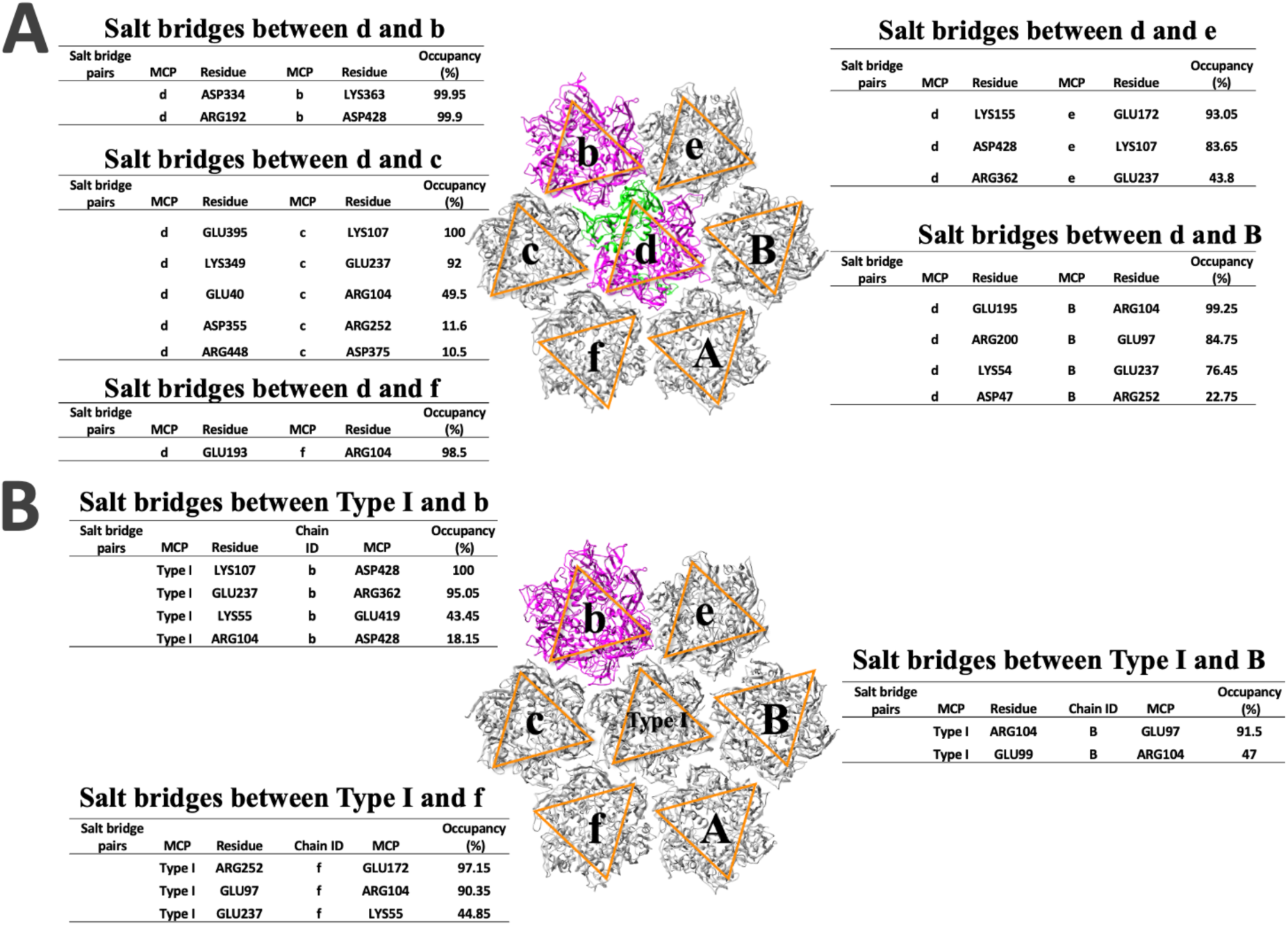
The salt bridges between Type Ⅴ/ Type I capsomer and surrounding capsomers. (A) Salt bridge pairs between Type V capsomer and surrounding capsomers. (B) Salt bridge pairs between Type I capsomer and surrounding capsomers. The salt bridges that rarely formed (<10% occupancy) in the simulations were ignored. Capsomers are labeled by orange triangles to show their orientations.

For the salt bridges shown in Figure 7A, several binding interfaces of the Type V capsomer exhibit high-occupancy salt bridges. The interface between the Type V capsomer and the c capsomer has the highest number of salt bridges, with five salt bridge pairs observed.

Remarkably, this interface also includes salt bridges with maximum occupancy (100%), indicating exceptionally strong and stable interactions. The interface between the Type V capsomer and the b capsomer features two salt bridges with nearly perfect occupancies (>99.90%). Additionally, high-occupancy salt bridges are present at the interfaces between the Type V capsomer and both the B capsomer and the e capsomer. Type V capsomer demonstrated a greater number of high-occupancy salt bridges, suggesting a stronger and more stable electrostatic network at its binding interface. These results align with the electrostatic potential and force analyses, which indicate that Type V capsomer contributes to a more robust local capsid microenvironment compared to Type I capsomer. Type V capsomer may play a potentially pivotal role in the early stages of viral capsid assembly due to its enhanced electrostatic interactions.

Moreover, Type V capsomer includes two MCPv1 monomers and one MCPv2 monomers. Most of the observed salt bridges involve the MCPv2 monomer or one of the MCPv1 monomers of the Type V pseudo-hexameric capsomer interacting with surrounding capsomers. Interestingly, although the other MCPv1 monomer of the Type V capsomer has the same sequence, it exhibits significantly weaker interactions with the surrounding capsomers. This disparity in interaction strengths by the two MCPv1 monomers suggests that the assembly order and the specific orientation of the Type V capsomer components are important factors in the assembly process.

The detailed mechanism of how the MCPv1 and MCPv2 monomers are assembled to a Type V capsomer will be in our future investigations.

In contrast, very few salt bridges were observed at the interfaces between the Type I capsomer and surrounding capsomers. This lack of stable interactions, combined with the electrostatic features described earlier, emphasis the significant role of the Type V capsomer in viral capsid assembly. The non-uniform distribution of Type V capsomers near the distinctive vertex of PBCV-1, although not fully understood, likely plays a pivotal role in the initiation of viral capsid assembly. The presence of charged Type V capsomers, along with the non-uniform distribution of Type II capsomers, suggests a coordinated mechanism that may regulate the assembly order of the viral capsid. Understanding this process could provide valuable insights into the molecular underpinnings of viral assembly and stability. Overall, the analysis of salt bridges in conjunction with electrostatic features highlights the critical role of the Type V capsomer in the assembly and stability of the PBCV-1 capsid. The charged nature and interaction patterns of the Type V capsomer suggest that it is a key component in the early stages of viral assembly, setting the stage for the formation of a stable and functional capsid.

## Conclusions

Type I capsomeres are the dominant type in the giant virus PBCV-1 capsid. However, some variants capsomers, such as Type Vs, exist at certain locations near the unique vertex, which may have some special functions. To study the function of the Type V capsomer, we replaced a Type V capsomer by a Type I capsomer to compare their roles with surrounding capsomers in PBCV-1. Through the combination of molecular dynamics (MD) simulations and electrostatic calculation tools such as DelPhi and DelPhiForce, we investigated the local capsid microenvironments induced by the Type V capsomer. This study aimed to elucidate the distinct electrostatic properties of the Type V/Type I pseudo-hexameric capsomers in the context of their surrounding environment and to identify the salt bridges that mediate interactions among PBCV-1 capsomers. Our findings reveal significant differences in the electrostatic features between the Type V and Type I capsomers when interacting with surrounding capsomers. Electrostatic potential calculations revealed distinct charge distributions, with the Type V capsomer inducing stronger and more attractive interactions, leading to a more stable capsid structure. Electric field line analysis supported these findings, showing denser fields and enhanced interaction strength for the Type V capsomer. Electrostatic force calculations further quantified the Type V capsomer’s greater attractive forces, emphasizing its critical role in capsid stability. Additionally, molecular dynamics simulations identified key interfacial residues forming stable salt bridges between the Type V capsomer with its surrounding capsomers, highlighting their importance in mediating capsomer interactions and maintaining structural integrity.

Overall, our study highlights the critical role of electrostatic interactions in the assembly of the PBCV-1 capsid, guiding capsomers to interact at optimal distances and orientations to ensure stable assembly. The presence of charged capsomers with a non-uniform distribution at key locations, such as near the unique vertex of the capsid, may be closely linked to the initiation of viral capsid assembly. These findings provide valuable insights into the molecular mechanisms that govern the assembly and stability of giant virus capsids and highlight the importance of considering electrostatic properties in understanding viral architecture. Future research will focus on further elucidating the detailed assembly mechanism of the Type V capsomer, including the order of assembly and conformational changes that occur during the formation of the viral capsid. Understanding these processes at a molecular level may offer new strategies for targeting and disrupting the assembly of viruses, contributing to the development of antiviral therapies.

## Acknowledgments

Research reported in this publication was supported by National Institute of General Medical Sciences (NIGMS) and National Institutes on Minority Health and Health Disparities (NIMHD) of the National Institutes of Health (NIH) under award number: SC1GM132043, R01GM129525, and U54MD007592. WG, EA, and CX also received support from Welch Foundation under Grant Number AH-2126-20220331.

