## Supplementary material for "Local Microenvironments of capsomer variants in the PBCV-1": SI

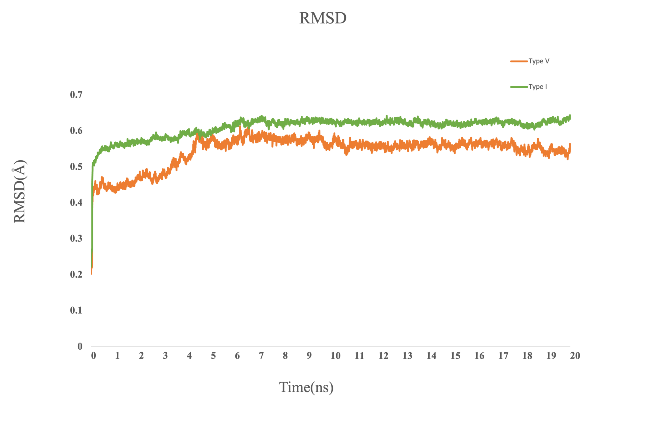


**Figure S1.** The RMSD of the simulations. The simulation results in RMSD vs Time (ns) of type V capsomers on giant virus PBCV-1 were represented with an orange line while the results of type I capsomers were displayed in green.
